# Molecular underpinnings of programming by early-life stress and the protective effects of early dietary ω6/ω3 ratio, basally and in response to LPS: hippocampal mRNA-miRNAs integrated approach

**DOI:** 10.1101/2022.08.06.503026

**Authors:** Kitty Reemst, Nicola Lopizzo, Maralinde R. Abbink, Hendrik J. Engelenburg, Annamaria Cattaneo, Aniko Korosi

## Abstract

Early-life stress (ELS) exposure increases the risk for mental disorders, including cognitive impairments later in life. We have previously demonstrated that a dietary low ω6/ω3 polyunsaturated fatty acid (PUFA) ratio protects against ELS-induced cognitive impairments. Several studies have implicated the neuroimmune system in the ELS and diet mediated effects, but currently the molecular pathways via which ELS and early diet exert their long-term impact are not yet fully understood. Here we study the effects of ELS and dietary PUFA ratio on hippocampal mRNA and miRNA expression in adulthood, both under basal and inflammatory conditions.

Male mice were exposed to chronic ELS by the limiting bedding and nesting material paradigm from postnatal day(P)2 to P9, and provided with a diet containing a high (15:1.1) or low (1.1:1) ω6 linoleic acid to ω3 alpha-linolenic acid ratio from P2 to P42. At P120, memory was assessed using the object location task. Subsequently, a single lipopolysaccharide (LPS) injection was given and 24 hours later hippocampal genome-wide mRNA and microRNA (miRNA) expression was measured using microarray.

An integrated miRNA – mRNA analysis revealed that ELS and early diet induced miRNA driven mRNA expression changes into adulthood. Under basal conditions both ELS and the diet affected molecular pathways related to hippocampal plasticity, with a low ω6/ω3 ratio diet specific activation of molecular pathways associated with improved hippocampal plasticity and learning and memory in mice previously exposed to ELS (e.g., CREB signaling and endocannabinoid neuronal synapse pathway). LPS induced miRNA and mRNA expression was strongly influenced by both ELS and early diet. In mice fed the high ω6/ω3 diet, LPS increased miRNA expression leading to activation of inflammatory pathways. In contrast, in mice fed the low ω6/ω3 diet, LPS reduced miRNA expression and altered target mRNA expression ultimately leading to limited activation of inflammatory signaling pathways and inhibition of pathways associated with hippocampal plasticity which was especially apparent in mice previously exposed to ELS.

This data provides molecular insights into how the low ω6/ω3 diet during development could exert its long-lasting beneficial effects on hippocampal plasticity and learning and memory especially in a vulnerable population exposed to stress early in life, providing the basis for the development of intervention strategies.

## 1. Introduction

There is ample evidence that early-life stress (ELS) is associated with increased risk for mental health problems, including cognitive impairments and alterations of brain structure and various forms of brain plasticity both from human^1–3^ as well as from animal studies^4–7^. Furthermore ELS leads to altered peripheral and central immune processes later in life^8–13^. Rodent studies have demonstrated that ELS leads to an aggravated neuroinflammatory reaction to later-life “secondary challenges” such as lipopolysaccharides (LPS)^14–17^ or accumulation of amyloid in the context of Alzheimer’s disease^13,18^. Such priming of the (neuro)immune system and exaggerated response to inflammatory challenges later in life has been proposed to contribute to the increased risk to develop psychopathology and cognitive impairment. Currently, the molecular underpinnings of these lasting effects are not fully understood and no intervention strategies are available.

We have proposed early-life nutrition to be a key player in mediating the ELS-induced effects as well as a potential target for intervention^19–24^. In particular adequate omega (ω)3 polyunsaturated fatty acids (PUFAs) levels during early-life have been acknowledged as an important determinant of later life mental health^25–27^. ω3 PUFAs are essential for normal brain development^28,29^, have a positive influence on cognition^30–32^ and have anti-inflammatory properties^33–36^.

Supporting the key role of PUFAs in early-life and the potential of early nutritional strategies for vulnerable populations exposed to ELS, we have recently reported that increasing the availability of ω3 PUFAs, via lowering the linoleic acid (LA)/α-linolenic acid (ALA) dietary ratio early in life (15:1 versus 1:1), protects against ELS-induced long-term cognitive dysfunction^20^. These beneficial effects of the diet on cognition were associated with a prevention of the ELS-induced changes in survival of hippocampal new-born neurons and microglial CD68 expression. In line with our findings, others have also reported dietary interventions with PUFA’s to be protective against ELS-induced changes in anxiety behaviours and cognitive functions in female rats^37^ and during adolescence^38^. Nonetheless currently, the molecular mechanisms underlying the protective effects of these diets are not yet understood.

In order to further our insights into the molecular pathways involved in both the long-term effects of ELS as well as the beneficial effect of the diet, we studied the hippocampal mRNA and microRNA (miRNA) expression profile. To gain insights into the upstream regulation of gene expression alterations, we integrated the mRNA with the miRNA expression profiles. miRNAs are evolutionary conserved, small non-coding RNAs (20-22 nucleotides in length) that play an important role in the post-transcriptional regulation of gene expression^39^. Indeed epigenetic mechanisms, including miRNAs, have been implicated in the mediation of early environmental cues into adult behavioral outcomes^40,41^, in particular, also in the context of ELS^42,43^ and nutrition^44–46^. Notably, dysregulation of miRNAs has been shown in several brain disorders including neurodegenerative and mental disorders^40,47^, for which ELS is a predisposing factor. Indeed alterations in plasma levels of miRNAs have been demonstrated in humans exposed to early life trauma^48,49^. While concerning pre-clinical evidence, there is quite some evidence from rodent studies demonstrating the role of brain miRNA expression in the effects of prenatal stress on brain and behaviour^48–52^. So far only few studies have explored the relationship between early postnatal stress and miRNAs^53–55^. For example, alterations in several brain miRNAs (medial prefrontal cortex, striatum and nucleus accumbens) were reported in ELS exposed rodents^53,54,56^, which were additionally modified after chronic stress exposure in adulthood^53,54^. Furthermore, there are few studies showing effects of fatty acids on miRNAs in peripheral tissues or cell-lines^45,57^ but none yet reported in the brain.

Thus we set out to investigate the long-term effects of ELS, on the integrated genome wide mRNA and miRNA expression profile in the hippocampus, assessed how these are impacted by dietary PUFA’s and studied these under basal conditions as well as in response to an inflammatory challenge in adulthood, as such a “secondary challenge” might be essential to unmask possible latent effects of ELS and the diet^13,58–62^.

## 2. Material and Methods

### 2.1 Animals

All mice (C57BI/6J) were kept under standard housing conditions with a temperature between 20 and 22°C, a 40 to 60% humidity level, a standard 12/12h light/dark schedule (lights on at 8AM), and provided with chow and water *ad libitum*. All experimental procedures were conducted under national law and European Union directives on animal experiments, and were approved by the animal welfare body of the University of Amsterdam.

Briefly, male mice were exposed to early-life stress (ELS) via limited bedding and nesting paradigm (postnatal day (P)2 to P9) (*paragraph 2.3*) and to an early diet (P2 – P42) with either high (15:1) or low (1.1:1) ω6 linoleic acid to ω3 alpha-linolenic acid ratio (*paragraph 2.4*). In adulthood mice were injected with either saline (SAL) or lipopolysaccharide (LPS) (*paragraph 2.5*). Hippocampal miRNA and mRNA were analysed using microarray (*section 2.8 and 2.9*) (**Fig. 1A**). The experimental groups were the following: control (CTL) mice fed a diet with high ω6/ω3 ratio and injected with saline: CTL-HRD-SAL; ELS exposed mice fed high ω6/ω3 ratio and injected with saline: ELS-HRD-SAL; control mice fed a diet with low ω6/ω3 ratio and injected with saline: CTL-LRD-SAL and ELS mice fed a diet with low ω6/ω3 ratio and injected with saline: ELS-LRD-SAL; control mice fed a diet with high ω6/ω3 ratio and injected with LPS: CTL-HRD-LPS; ELS exposed mice fed high ω6/ω3 ratio and injected with LPS: ELS-HRD-LPS; control mice fed a diet with low ω6/ω3 ratio and injected with LPS: CTL-LRD-LPS and ELS mice fed a diet with low ω6/ω3 ratio and injected with LPS: ELS-LRD-LPS. The sample size per experimental group was 8 for the mRNA and miRNA expression analysis.

**Figure 1.**
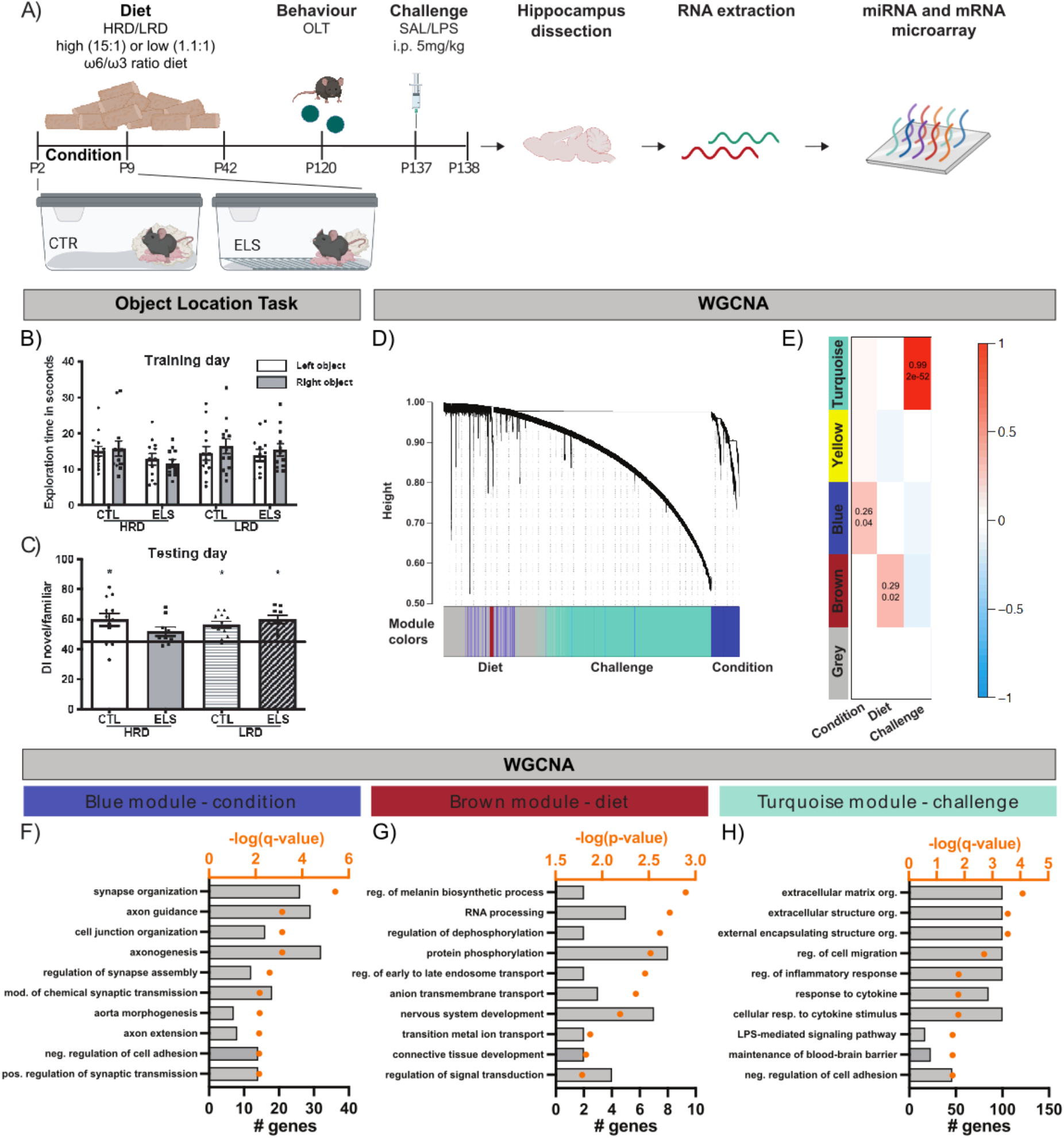
Early low ω6/ω3 ratio protects against ELS induced cognitive deficits and WGCNA shows gene co-expression modules related to condition, diet and challenge. **A)** Experimental timeline generated with Biorender.com. **B)** Object exploration during training day of OLT is not affected by condition or diet. **C)** All experimental groups learn significantly better than chance level (t-test against 45, p<0.05), except ELS mice fed the HRD. **D)** Detected gene co-expression modules by WCGNA. **E)** Pearson correlation of modules detected with WGCNA and predictor variables (Condition (CTL/ELS), Diet (HRD/LRD), Challenge (SAL/LPS), p-values is indicated as number and R2 number for significant (p<0.05) and as color for all correlations. **FGH)** Top 10 significantly enriched GO terms associated with the blue (**F**; condition; q-value < 0.05), brown (**G**; diet; p-value < 0.05) and turquoise (**H**; challenge; q-value < 0.05) module genes. Abbreviations: CTL: control, ELS: early-life stress, SAL: saline, LPS: lipopolysaccharide, HRD: high ω6/ω3 ratio diet, LRD: low ω6/ω3 ratio diet, WGCNA: weighted gene co-expression network analysis, GO: gene ontology, reg.: regulation, mod.: modification, neg.: negative, pos.: positive.

Data on bodyweight, food intake and plasma measurements can be found in Reemst et al (2022)^63^.

### 2.2 Breeding

Experimental mice were bred in house to standardize the perinatal environment. 10-week-old female and 8-week-old male mice were purchased from Harlan Laboratories B.V. (Venray, The Netherlands) and habituated for two weeks before onset of breeding. After the habituation period, two females and one male were housed together for one week to allow mating. Breeding males were removed after one week and after another week of paired-housing, pregnant primiparous females were individually housed in a standard cage with a filtertop. To ensure a stable, quiet environment, cages were placed in a ventilated, airflow-controlled cabinet. Birth of pups was monitored every 24 hours. Litters born before 9:00 AM were considered postnatal day (P)0 on the previous day.

### 2.3 Early-life stress paradigm

Chronic ELS was induced via using the limited bedding and nesting material (LBN) paradigm from P2 to P9 as described previously by our group and others^4,13,20,64^. On the morning of P2, dams and pups were randomly assigned to the CTL or ELS condition. Litters were culled to six pups with a minimum of 5 pups to prevent maternal care variation due to variable litter size. Litters included at least one male and one female. At P2, dams and pups were weighed and housed under CTL or ELS conditions. CTL cages contained standard amounts of sawdust bedding and one square, cotton piece of nesting material (5×5 cm; Technilab-BMI, Someren, The Netherlands). ELS cages contained fewer amounts of sawdust bedding, only covering the bottom of the cage, a fine-gauge stainless steel mesh raised 1 cm above the cage floor, and half a square cotton piece of nesting material (2,5×5 cm). Cages were covered with a filtertop and placed in a ventilated, airflow-controlled cabinet to ensure a stable, quiet environment and reduce external stressors. Throughout all procedures, manipulation was kept to a minimum to avoid handling effects and mice were left undisturbed until P9. On the morning of P9, pups were weighed and moved to standard cages. Mice were weaned at P21 and housed with same-sex littermates in groups of 2 to 3 mice per cage. Only male offspring was used for experimental procedures (the experimental timeline is represented in **figure 1A**).

### 2.4 Experimental diets

Dams were assigned to the American Institute of Nutrition-93 (AIN-93G/M) semi-synthetic diet throughout the breeding period^65^. Experimental diets were provided from P2 onwards to dam with litter, and after weaning (P21) offspring were kept on their respective diet until P42. The two experimental diets (Ssniff-Spezialdiäten GmbH, Soest, Germany) were semi-synthetic differing only in ω6 linoleic (LA)/ω3 α-linolenic (ALA) ratio that was either a high (15:1) or low (1.1:1). The diets were isocaloric and contained a macro- and micronutrient composition according to the AIN93-G purified diets for laboratory rodents^65^ (**supplementary table S1**). Following dietary intervention at P42, all mice were fed AIN-93M until the end of the experiment.

### 2.5 Behavioral testing

At P120, 14 mice of each of the 4 experimental groups were tested in the object location task (OLT). In order to perform all behavioral testing in the active phase, the light-dark cycle was reversed (reversed 12/12h light/dark schedule, lights off at 8AM) 4 weeks before onset of testing. Behavior tests were recorded by Ethovision (Noldus, The Netherlands) and scored manually using Observer software (Noldus) by one experimenter who was blind to the conditions. Prior to the OLT, mice were handled for 3 days to diminish possible stress induced by the experimenter. During the habituation phase, mice were allowed to explore the testing arena (24×31×27 cm box covered with a small amount of sawdust) for 5 min on 3 subsequent days. On the training day, two identical objects placed equidistant from each other and from the wall of the arena were placed in the testing arena and mice could explore the objects for 5 minutes. On the testing day (24 hours later), one of the objects was relocated (the object and direction of relocation was randomly assigned) within the arena and again mice were allowed to explore the objects for 5 minutes. For all days, boxes and objects were cleaned with 25% ethanol after each tested animal. Exploration was defined as mice touching the object with their nose. Mice were excluded from the task when a preference for one of the objects was present during the training phase, or when the total exploration time in either the training or testing phase was below 10 seconds. Cognitive performance was assessed using the discrimination index (DI) of the testing day: the time spent exploring the novel object divided by total exploration time of both objects.

### 2.6 Lipopolysaccharide injection

Approximately one week (5-9 days) after the end of the behavioral test, mice were weighed and received an intraperitoneal (i.p.) injection of sterile saline (SAL) or 5 mg/kg lipopolysaccharide (LPS, strain O111:B4, Sigma-Aldrich) dissolved in sterile saline^14,66^. 24 hours after LPS injection, mice were weighted and sacrificed via rapid decapitation. Hippocampi were extracted and stored at -80°C until further processing.

### 2.7 Statistical analyses for behaviour

Data were analyzed using SPSS 20.0 (IBM software) and Graphpad Prism 5 (Graphpad software). Data were expressed as mean ± standard error of the mean (SEM) and considered statistically significant when p<0.05. Cognitive performance in the OLT was assessed using one-sample t-test against 50% and two-way-ANOVA. In case of significant interaction effects, post hoc analyses were performed using Tukey’s post hoc test. As multiple mice from a litter were included in experiments, litter corrections were performed when a significant contribution of litter was found in a mixed model analysis with litter included as random factor.

### 2.8 RNA isolation and mRNA/miRNAs microarray analyses

Hippocampal RNA of 8 mice per experimental group was extracted using the TRIzol method (TRIzol Invitrogen) followed by application of RNA Clean & concentrator according to the manufacturer’s instructions (Clean and concentrator -25, Zymo Research).

#### 2.8.1 mRNA microarray analyses

A total amount of two nanogram of total RNA was amplified using the GeneChip Pico Reagent Kit (Thermo Fisher Scientific) generating biotinylated double-stranded cDNA. The labeled samples were hybridized to mouse Clariom S array plate (Thermo Fisher Scientific). Washing, staining, and scanning was performed using the GeneTitan Wash and Stain Kit for WT Array Plates, and the GeneTitan Instrument (Thermo Fisher Scientific) performed by the MicroArray Department (MAD, University of Amsterdam, The Netherlands).

#### 2.8.2 microRNA microarray analyses

A total amount of 250 nanogram of total RNA from each sample was processed with the FlashTag Biotin HSR RNA Labeling kit (Thermofisher, Waltham, MA, USA) and subsequently hybridized onto the GeneChip miRNA 4.1 Arrays (Thermofisher, Waltham, MA, USA), on a GeneAtlas platform (Affymetrix, Santa Clara, CA, USA). The GeneChip miRNA 4.1 Array Strip shows a comprehensive coverage as they are designed to interrogate all mature miRNA sequences in miRBase Release 20 (online miRNA database, http://www.mirbase.org). Washing/staining and scanning procedures were respectively conducted on the Fluidics Station and the GeneChip Scanner of a GeneAtlas instrument (Affymetrix, Santa Clara, CA, USA) following the manufacturer’s instructions.

### 2.9 Strategy and downstream gene and microRNA expression analysis

Our data analysis strategy was as follows. First, we performed an unsupervised Weighted Gene Co-expression Network Analysis (WGCNA) on gene expression of all of our samples without specifying experimental groups, in order to identify modules of genes with similar expression patterns in an unbiased manner. The genes making up modules that correlated with our predictor variables (*ELS, diet, challenge*) were analyzed by gene ontology in order to learn more about the biological processes they are involved with. Next, we focused our analysis on the integrated analysis of miRNA and mRNA in order to understand in which ways miRNAs driven mRNA expression changes are involved in the effects of ELS and early diet, both under basal conditions as in response to LPS. Ingenuity Pathway Analysis (IPA) was performed on the differentially expressed target mRNAs affected by differentially expressed miRNAs, to learn more about the associated molecular pathways. The independent mRNA and miRNA expression profiles are clearly also of great interest however describing those in detail is out of the scope of this paper. Therefore, we do not report them in our main result section, but lists with differentially expressed mRNAs and miRNAs of the considered contrasts can be found in **Supplementary table S2 and S3** respectively.

#### 2.9.1 Weighted Gene Co-expression Network Analysis and gene ontology analysis

WGCNA was applied on gene expression data using the WGCNA package (v1. 70-3, Langfelder & Horvath, 2008). The 8000 genes with most variation in gene expression over all samples were selected. An unsigned topological overlap matrix (TOM) was constructed using a power adjacency function with a soft power threshold of 4. Subsequently, a dendrogram was constructed using average linkage hierarchical clustering. Modules were created with a dynamic tree cut of the dendrogram (Langfelder et al 2008), using a recommended deep-split of 2 (Langfelder, 2015), minimum size of 20 genes, minimum eigengene connectivity (kME) of 0.3 and modules with height < 0.3 were merged. Metadata variables (condition, diet, treatment) were converted to binary variables and were correlated (Pearson correlation) to the previously identified module eigengenes. A correlation with p-value<0.05 was considered significant.

Gene ontology (GO) analysis for differentially expressed genes and gene modules was performed with enrichR (v3.0, Xie et al., 2021). The database “GO_Biological_Process_2021” was used to identify enriched GO terms and GOs were accounted as significantly enriched with an FDR adjusted p-value<0.05. In cases where no GO terms were enriched with an adjusted p-value<0.05, GO terms with a p-value<0.05 were reported.

#### 2.9.2 Integrated miRNA/mRNA analysis

Expression of raw data was imported and analyzed with the software Partek Genomic Suite 6.6 (Partek, St. Louis, MO, USA). All samples passed the criteria for hybridization controls, labelling controls and 3′/5′ Metrics. Background correction was conducted using Robust Multi-strip Average (RMA) (Irizarry et al., 2003) to remove noise from auto fluorescence. After background correction, normalization was conducted using Quantiles normalization (Bolstad et al., 2003) to normalize the distribution of probe intensities among different microarray chips. Subsequently, a summarization step was conducted using a linear median polish algorithm (Tukey, 1977) to integrate probe intensities to compute the expression levels for each mRNA transcript. Principal-component analysis (PCA) was carried out to identify possible outliers and major effects in the data. No significant outliers were observed; however, a batch effect was detected in both the mRNA and miRNA datasets, which was corrected using the relative “Remove batch effect” option in Partek Genomics Suite. After quality control of the data, linear contrasts were performed for several contrasts (Table 1) identifying differentially expressed mRNAs and miRNAs. Fold change (FC) > |1.2| and p-value <0.05 were regarded as significant.

**Table 1.** Differentially expressed miRNAs per contrast from integrated miRNA-mRNA analysis. p-value < 0.05 and fold change > |1.2| (see supplementary table S5 for specifics per miRNA).

| Contrast | Upregulated in A versus B | % | Downregulated in A versus B | % |
| --- | --- | --- | --- | --- |
| **Diet effect in control mice**<br>*SAL-CTL: LRD vs HRD* | miR-30b-5p, miR-671-5p, miR-7a-5p, miR-23a-3p, miR-323-3p, miR-6516-5p, miR-324-3p, miR-378a-3p, miR-411-5p, miR-370-3p, miR-204-5p, miR-25-3p, miR-504-3p, miR-27b-3p, miR-410-3p, miR-384-5p, miR-29c-3p, miR-127-5p, miR-495-3p, miR-331-3p, miR-874-3p, miR-9-5p, miR-30a-5p, miR-874-5p, miR-153-3p, miR-381-3p, miR-431-5p, miR-15a-5p | 90 | miR-200c-3p, miR-5107-5p, miR-3072-5p | 10 |
| **ELS effect in mice fed the HRD**<br>*SAL-HRD: ELS vs CTL* | – | 0 | miR-200c-3p, miR-1224-5p, miR-183-5p, miR-7018-5p, miR-182-5p, miR-6906-5p, miR-6954-5p, miR-6939-5p | 100 |
| **ELS effect in mice fed the LRD**<br>*SAL-LRD: ELS vs CTL* | miR-346-5p, miR-338-5p, miR-429-3p, miR-7020-5p | 40 | miR-7a-5p, miR-7220-5p, miR-29c-3p, miR-30a-5p, miR-195a-5p, miR-19b-3p | 60 |
| **LPS effect in CTL mice fed the HRD**<br>*CTL-HRD: LPS vs SAL* | miR-6968-5p, miR-92a-3p, miR-21a-3p, miR-30c-1-3p, miR-323-3p, miR-23b-5p, miR-1982-5p, miR-128-1-5p, miR-6945-5p, miR-128-2-5p, miR-296-5p, miR-6911-5p, miR-20a-5p, miR-135a-2-3p, miR-29c-3p, miR-7008-5p, miR-380-5p, miR-1968-5p, miR-346-3p, miR-122-5p, miR-7b-5p | 72 | miR-3072-5p, miR-1894-5p, miR-7018-5p, miR-3064-5p, miR-3474, miR-470-5p, miR-3966, miR-503-5p | 28 |
| **LPS effect in CTL mice fed the LRD**<br>*CTL-LRD: LPS vs SAL* | miR-21a-5p, miR-669h-3p | 10 | miR-320-3p, miR-671-5p, miR-7665-5p, miR-500-3p, miR-23a-3p, miR-6516-5p, miR-150-5p, miR-350-3p, miR-212-5p, miR-1231-5p, miR-7235-5p, miR-204-5p, miR-143-3p, miR-410-3p, miR-7028-5p, miR-6934-5p, miR-7038-3p, miR-329-3p | 90 |
| **LPS effect in ELS mice fed the HRD**<br>*ELS-HRD: LPS vs SAL* | miR-762, miR-6968-5p, miR-485-5p, miR-342-3p, miR-320-3p, miR-668-3p, miR-92a-3p, miR-129-2-3p, miR-23a-3p, miR-27b-5p, miR-7215-3p, miR-744-5p, miR-138-2-3p, miR-99b-5p, miR-1982-5p, miR-346-5p, miR-191-5p, miR-187-3p, miR-128-2-5p, miR-1224-5p, miR-370-3p, miR-138-5p, miR-127-3p, miR-18a-5p, miR-151-5p, miR-92b-3p, miR-135a-2-3p, miR-181a-5p, miR-7081-5p, miR-331-3p, miR-146a-5p, miR-7119-3p, miR-7648-3p, miR-3473b, miR-501-3p, miR-3473e | 95 | miR-3059-5p, miR-1962 | 5 |
| **LPS effect in ELS mice fed the LRD**<br>*ELS-LRD: LPS vs SAL* | miR-6968-5p, miR-211-3p, miR-5110, miR-365-1-5p, miR-296-5p, miR-6953-5p, miR-297a-5p, miR-7052-5p, miR-7216-5p, miR-1956 | 10 | miR-130a-3p, miR-378d, miR-342-3p, miR-320-3p, miR-6987-5p, miR-30b-5p, miR-129-2-3p, miR-23b-3p, miR-671-5p, miR-7665-5p, miR-3084-3p, miR-30c-1-3p, miR-500-3p, miR-23a-3p, miR-543-3p, miR-323-3p, miR-6516-5p, miR-873a-3p, miR-539-5p, miR-181d-5p, miR-30e-3p, miR-669a-5p, miR-669p-5p, miR-185-3p, miR-378b, miR-138-2-3p, miR-99b-5p, miR-339-5p, miR-99a-5p, miR-191-5p, miR-152-3p, miR-149-5p, miR-338-5p, miR-187-3p, miR-378a-3p, miR-325-5p, miR-150-5p, miR-28a-5p, miR-222-3p, miR-181c-5p, miR-1231-5p, miR-106a-5p, miR-370-3p, miR-138-5p, miR-1843a-5p, miR-328-3p, miR-107-3p, miR-127-3p, miR-181b-5p, miR-17-3p, miR-134-5p, miR-221-3p, miR-28a-3p, miR-6769b-5p, miR-423-3p, miR-25-3p, miR-26b-5p, miR-143-3p, miR-125a-5p, miR-125b-5p, miR-412-5p, miR-27b-3p, miR-690, miR-139-5p, miR-410-3p, miR-421-3p, miR-384-5p, miR-181a-5p, miR-145a-5p, miR-532-5p, miR-425-5p, miR-708-5p, miR-30b-3p, miR-495-3p, miR-103-3p, miR-20b-5p, miR-378c, miR-6972-5p, miR-376b-3p, miR-6965-5p, miR-423-5p, miR-351-5p, miR-504-5p, miR-409-5p, miR-193b-3p, miR-100-5p, miR-34b-3p, miR-129-5p, miR-494-3p, miR-7684-5p | 90 |

As mentioned above, in this paper we focused our analysis on the combined actions of differentially expressed mRNAs and miRNAs in our dataset. Using a specific sub-feature “combine” in Partek Genomic Suite 6.6, we integrated differentially expressed miRNAs with their predicted mRNA targets from the list of differentially expressed mRNAs. List of significant target mRNAs were subjected to pathway analyses by using Ingenuity Pathway Analyses Software (IPA, Ingenuity System Inc, USA http://www.ingenuity.com). The “Core Analysis” function included in IPA was used to understand the data in the context of biological processes, pathways, networks and upstream regulators associated with each condition of interest.

Two metrics were used to identify the most important downstream effects of differentially expressed mRNAs: p-value and activation z-score. The p-value, calculated with the Fischer’s exact test, indicates the likelihood that the association between a set of genes in our dataset and a biological function is significant, with a threshold of 0.05 (^10^log(p-value) = 1.3). The activation z-score was used to infer likely activation states of biological functions based on comparison with an IPA model that assigns random regulation directions. A positive or negative z-score indicates increased or decreased functional activity in comparison A versus B. Pathways/biological processes with a z-score ≥ |2| were regarded to be significantly activated or inhibited.

### 2.10 General strategy

In order to tackle the molecular substrates of the long-term effect of ELS and those of the beneficial effect of early dietary PUFAs we exposed mice to CTL or ELS condition. Half of the mice were fed early in life the HRD, a diet with a similar fatty acid ratio as in a standard rodent diet^65^ and under which ELS mice recapitulated the expected ELS-induced deficits in cognition and brain plasticity^20^. The other half received the LRD, which we have earlier demonstrated to protect against the ELS-induced deficits^20^. In adulthood mice were exposed to either LPS or saline (SAL) to unmask potential latent effects of ELS. Thus, we have three predictor variables: condition (CTL/ELS), diet (HRD/LRD) and challenge (SAL/LPS), leading to eight experimental groups: CTL-HRD-SAL, ELS-HRD-SAL, CTL-LRD-SAL, ELS-LRD-SAL, CTL-HRD-LPS, ELS-HRD-LPS, CTL-LRD-LPS, ELS-LRD-LPS. Considering the complexity of this design, in order to describe and disentangle the physiological effects and molecular underpinnings of ELS and the protective effects of the diet, we started by studying the long-term effects of the two different dietary PUFA ratios in CTL mice. While this is not the primary question in our study, it is key to understand the long-term effects of the diet at baseline in CTL mice to be able to dissect the specific effects of ELS under the different dietary exposures. To understand the molecular substrates of the ELS-induced deficits we studied the effects of ELS in mice fed the HRD as compared to their CTL-HRD counterpart. Thereafter, to gain further understanding of the molecular pathways underlying the beneficial effect of the LRD we studied the impact of ELS in mice fed the LRD as compared their CTL-LRD counterpart and assessed if these effects differed from those detected in mice (CTL and ELS) fed HRD. Lastly, we studied how the early-life environment (ELS and diet) impacted the expression profiles in response to LPS.

## 3. Results

### 3.1 Dietary intervention with low ω6/ω3 ratio from P2 – P42 prevents ELS-induced cognitive impairments in the object location task

Mice underwent object location task (OLT) at P120. No differences in total exploration time between groups were observed during the training phase (**Fig. 1B**). In the testing phase, CTL mice on high and low ω-6/ω-3 diet explored the relocated, novel object more than the familiar object (**Fig. 1C**; one-sample t-test CTL-HRD: t_8_=2.322, p=0.041; CTL-LRD t_11_=2.999, p=0.012). ELS mice fed a HRD showed impaired object location memory, which was not the case in ELS mice fed the LRD (**Fig. 1C**; one-sample t-test ELS-HRD: t_8_=0.595, p=0.5675; ELS-LRD t_8_=3.771 p=0.0055). Analysis of OLT performance by two-way ANOVA revealed no overall differences between groups.

### 3.2 Effects of early-life stress, dietary ω6/ω3 ratio on hippocampal mRNA and miRNA expression profile under basal conditions and in response to LPS

First, we will describe the unsupervised analyses of gene expression data by Weighted Gene Co-expression Network Analysis (WGCNA), followed by the integrated analysis of differentially expressed mRNAs and miRNAs.

#### 3.2.1 WGCNA

To identify modules of genes with similar expression patterns in an unbiased manner, WGCNA was performed on all gene expression data. Five co-expression modules were identified of which three significantly correlated with one of the predictor variables (**Fig. 1D**). One module significantly correlated with *condition* (blue, R^2^= 0.26, p<0.05), one with *diet* (brown, R^2^= 0.29, p<0.05) and one with *challenge* (turquoise, R^2^= 0.99, p<0.001) (**Fig. 1E**). Gene Ontology analysis indicated that the blue module genes (*condition*) are involved in structural and functional components of the synapse, axonogenesis and cell-cell communication (**Fig. 1F**). For the brown module (*diet*), genes are associated with protein phosphorylation and nervous system development (**Fig. 1G**). The turquoise module genes (*challenge*) are involved in extracellular matrix organization and the inflammatory response (**Fig. 1H**). The top 10 co-expression module-genes with highest centrality (*hub-genes*) can be found in **supplementary table S4**.

#### 3.2.2 Effects of ELS and early dietary PUFA ratio on integrated gene and miRNA expression profiles under basal conditions

Detailed results of the integrated miRNA-mRNA analysis (all differentially expressed mRNA, miRNAs, p-values and fold-changes per contrast) can be found in **supplementary table S5**. Additionally, an overview of the impacted miRNAs in the various contrast is shown in **table 1**. As mentioned above, we first studied the long-term impact of early dietary PUFA ratio on hippocampal mRNA and miRNA expression under basal conditions. To determine this effect, CTL mice fed either the HRD or LRD were compared (CTL-SAL: LRD versus HRD). 71 mRNAs were detected, targeted by 31 miRNAs. Notably, the majority of miRNAs were upregulated by the LRD (90%; e.g., miR-30b-5p, miR-7a-5p, miR-27b-3p, miR-29c-3p, miR-9-5p, miR-30a-5p) and three miRNAs were downregulated (10%; miR-200c-3p, miR-5107-5p, miR-3072-5p) (**Table 1**). IPA on the detected target mRNAs revealed involvement in cyclic adenosine monophosphate (cAMP)-response element binding protein (CREB) signaling in Neurons”, “Phagosome Formation”, and “Adipogenesis pathway” (**Table 2A**).

**Table 2.** Top 10 IPA pathways associated with the outcome mRNAs after integrated miRNA-mRNA analysis of diet and ELS effects. IPA: Ingenuity Pathway analysis, ELS: early-life stress, SAL: saline, LRD: low ω6/ω3 ratio diet, HRD: high ω6/ω3 ratio diet. p-value < 0.05, z-score > |2|

**A. Effect of diet on target mRNA expression as analyzed by IPA**
| Comparison | Ingenuity Canonical Pathways | p-value | zScore | #mRNAs | Molecules |
| --- | --- | --- | --- | --- | --- |
| **Diet effect in CTL mice**<br>*CTL-SAL: LRD vs HRD* | CREB Signaling in Neurons | 0.028 | 1.34 | 5 | BMPR1A, EDNRB, FZD7, GRM4, HGF |
|  | Phagosome Formation | 0.013 | 0.82 | 6 | EDNRB, FZD7, GRM4, MAPK4, PLA2G6, WIPF1 |
|  | Role of Osteoblasts, Osteoclasts and Chondrocytes in Rheumatoid Arthritis | 0.000 | — | 5 | BMPR1A, FZD7, LRP5, NFKBID, WNT10B |
|  | Adipogenesis pathway | 0.001 | — | 4 | BMPR1A, FZD7, PPARG, WNT10B |
|  | Factors Promoting Cardiogenesis in Vertebrates | 0.001 | — | 4 | BMPR1A, FZD7, LRP5, WNT10B |
|  | Methylthiopropionate Biosynthesis | 0.003 | — | 1 | ADI1 |
|  | MIF-mediated Glucocorticoid Regulation | 0.005 | — | 2 | NFKBID, PLA2G6 |
|  | Role of NANOG in Mammalian Embryonic Stem Cell Pluripotency | 0.005 | — | 3 | BMPR1A, FZD7, WNT10B |
|  | Hepatic Fibrosis Signaling Pathway | 0.007 | — | 5 | FZD7, LRP5, NFKBID, PPARG, WNT10B |
|  | MIF Regulation of Innate Immunity | 0.007 | — | 2 | NFKBID, PLA2G6 |

**B. Effect of ELS on target mRNA expression in mice fed the HRD or LRD as analyzed by IPA**
| Comparison | Ingenuity Canonical Pathways | p-value | zScore | #mRNAs | Molecules |
| --- | --- | --- | --- | --- | --- |
| **ELS effect in mice fed the HRD**<br>*HRD-SAL: ELS vs CTL* | Protein Kinase A Signaling | 0.009 | — | 3 | GNA13, PTPN13, PTPN14 |
|  | Ephrin Receptor Signaling | 0.019 | — | 2 | GNA13, PTPN13 |
|  | AMPK Signaling | 0.027 | — | 2 | GNA13, PPM1F |
|  | TWEAK Signaling | 0.039 | — | 1 | TRAF3 |
|  | A proliferation-inducing ligand (APRIL) Mediated Signaling | 0.044 | — | 1 | TRAF3 |
|  | B Cell Activating Factor Signaling | 0.045 | — | 1 | TRAF3 |
|  | G Protein Signaling Mediated by Tubby | 0.046 | — | 1 | GNA13 |
|  | IL-23 Signaling Pathway | 0.048 | — | 1 | RORA |
|  | Role of RIG1-like Receptors in Antiviral Innate Immunity | 0.048 | — | 1 | TRAF3 |
| **ELS effect in mice fed the LRD**<br>*LRD-SAL: ELS vs CTL* | Hepatic Fibrosis Signaling Pathway | 0.000 | 2.45 | 8 | CACNA1D, CACNA1E, CACNG4, FGFR1, FLT1, KDR, LRP1, PDGFRB |
|  | CREB Signaling in Neurons | 0.000 | 2.33 | 9 | CACNA1D, CACNA1E, CACNG4, FGFR1, FLT1, GRIN1, HGF, KDR, PDGFRB |
|  | Nitric Oxide Signaling in the Cardiovascular System | 0.000 | 2 | 5 | CACNA1D, CACNA1E, CACNG4, FLT1, KDR |
|  | IL-15 Production | 0.001 | 2 | 4 | FGFR1, FLT1, KDR, PDGFRB |
|  | Endocannabinoid Neuronal Synapse Pathway | 0.002 | 2 | 4 | CACNA1D, CACNA1E, CACNG4, GRIN1 |
|  | PTEN Signaling | 0.002 | -2 | 4 | FGFR1, FLT1, KDR, PDGFRB |
|  | Calcium Signaling | 0.008 | 2 | 4 | CACNA1D, CACNA1E, CACNG4, GRIN1 |
|  | Opioid Signaling Pathway | 0.018 | 2 | 4 | CACNA1D, CACNA1E, CACNG4, GRIN1 |
|  | White Adipose Tissue Browning Pathway | <0.000 | 1.63 | 6 | CACNA1D, CACNA1E, CACNG4, FGFR1, KDR, LIPE |
|  | STAT3 Pathway | 0.000 | 1.34 | 5 | FGFR1, FLT1, HGF, KDR, PDGFRB |

Next, we assessed the molecular pathways impacted by ELS effects in mice fed the standard HRD (HRD-SAL: ELS versus CTL). Integrated analysis revealed 27 mRNAs that were targeted by 8 miRNAs (**Fig. 2AB**). All detected miRNAs were downregulated (100%) by ELS such as for example miR-200c-3p and miR-182-5p (**Table 1)**. IPA on the target mRNA revealed involvement in “Protein Kinase A Signaling”, “Ephrin Receptor signaling” and “AMPK signaling” (**Table 2B**). In order to investigate whether the impact of condition on mRNA and miRNA profiles was dependent on the early diet, we compared data of CTL and ELS exposed mice fed either the HRD or LRD (HRD-SAL: ELS versus CTL and LRD-SAL: ELS versus CTL). Remarkably, most ELS-impacted mRNAs and miRNAs were unique depending on the diet and there were more ELS-induced DEGs and DEMs in mice fed the LRD (88 DEGs that were regulated by 10 DEMs; **Fig. 2A, B**). In particular, in ELS mice fed the LRD as compared to CTL mice fed the LRD, both upregulated (40%; e.g., miR-338-5p) and downregulated (60%; e.g., miR-7a-5p, miR-29c-3p, miR-30a-5p, miR-195a-5p) DEMs were detected. IPA on the 88 differentially expressed target mRNAs of these miRNAs revealed significant activation of pathways including for example “CREB signaling in Neurons”, “Nitric Oxide Signaling”, “IL15 production” and “Endocannabinoid Neuronal Synapse Pathway” (**Table 2B**; **Fig. 2C**).

**Figure 2.**
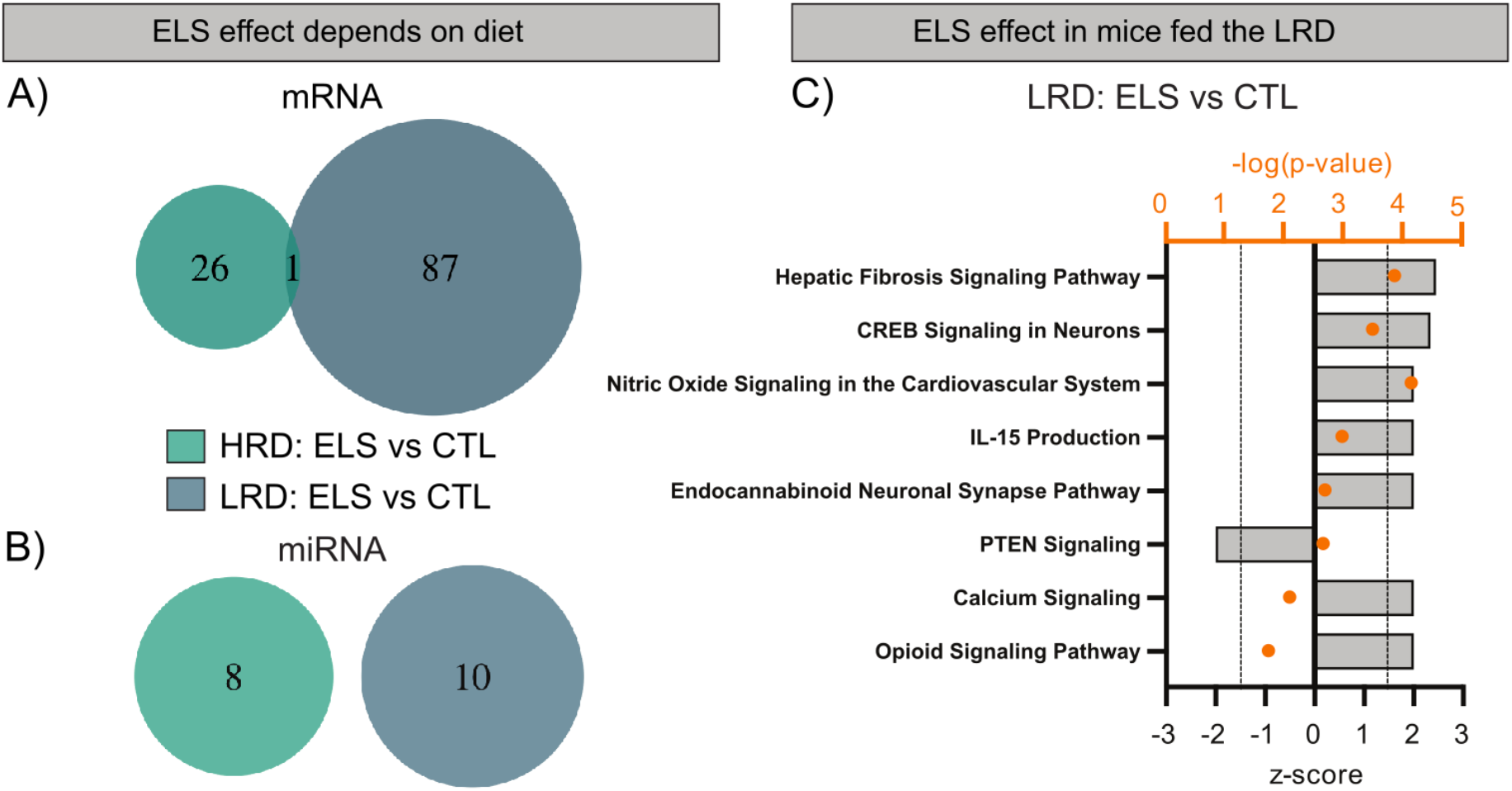
ELS effect on hippocampal miRNA and target mRNA expression depends on early dietary ω6/ω3 ratio. **AB)** ELS effect on hippocampal mRNA **(A)** and miRNA **(B)** expression depends on the early diet. **C)** IPA showing molecular pathways (p-value <0.05 and z-score > |2|) involved with the impact of ELS in mice fed the LRD. Abbreviations: CTL: control, ELS: early-life stress, HRD: high ω6/ω3 ratio diet, LRD: low ω6/ω3 ratio diet.

To summarize, the early dietary ω6/ω3 PUFA ratio affected several miRNAs and their target mRNAs in the adult hippocampus, with LRD specific increase in miRNA expression and the most prominent pathways that emerge were associated with neuronal plasticity processes such as CREB signaling and phagosome formation. The diet not only had a large impact on miRNAs and target mRNAs under control conditions, but also greatly impacted the ELS-induced effects. In fact, ELS (under the standard-HRD) reduced expression of several miRNAs and affected mRNAs associated with hippocampal PKA, ephrin and AMPK signaling. In contrast, when mice were fed an LRD early in life, ELS altered more miRNAs and target mRNAs, which led to an activation of pathways associated with hippocampal plasticity and learning and memory.

#### 3.2.3 Effects of ELS and early dietary PUFA ratio on integrated gene and miRNA expression profiles in response to LPS

To determine the impact of early dietary ω6/ω3 ratio on mRNA and miRNA expression in response to LPS we first studied this in CTL mice (*CTL-HRD: LPS versus SAL* and *CTL-LRD: LPS versus SAL*). Integrated analysis of differentially expressed mRNAs and miRNAs revealed that the impact of LPS in mice fed the HRD led to 614 mRNAs that were regulated by 29 miRNAs. In CTL mice fed the LRD, 580 mRNAs regulated by 20 miRNAs were detected. Notably, most of the mRNAs and all of the miRNAs were unique for each diet (**Fig. 3AB**), and remarkably while in mice fed the HRD most miRNAs were upregulated (72%; e.g. miR-27b-5p, miR-187-3p, miR-181a-5p, miR-146a-5p), in mice fed the LRD the majority was downregulated (90%; e.g. miR-212-5p, miR-204-5p) in response to LPS (**Table 1**; **supplementary table S5**). For the HRD, IPA on the target genes revealed LPS induced activation of pathways associated with immune functions and disease states such as “Hepatic Fibrosis Signaling Pathway”, “IL-6 signaling”, “Production of NO and reactive Oxygen Species (ROS) in macrophages”, “PPARα/RXRα activation” and TGFβ signaling” (**Fig. 3C; supplementary table S6**). For the LRD, none of the affected pathways were the same as affected by LPS in mice fed the HRD. LPS induced for example activation of “HIFa signaling” and “Signaling by Rho Family GTPases” and inhibition of “cAMP-signaling”, “CREB-signaling” and “Endocannabinoid pathway” (**Fig. 3D; supplementary table S6**).

**Figure 3.**
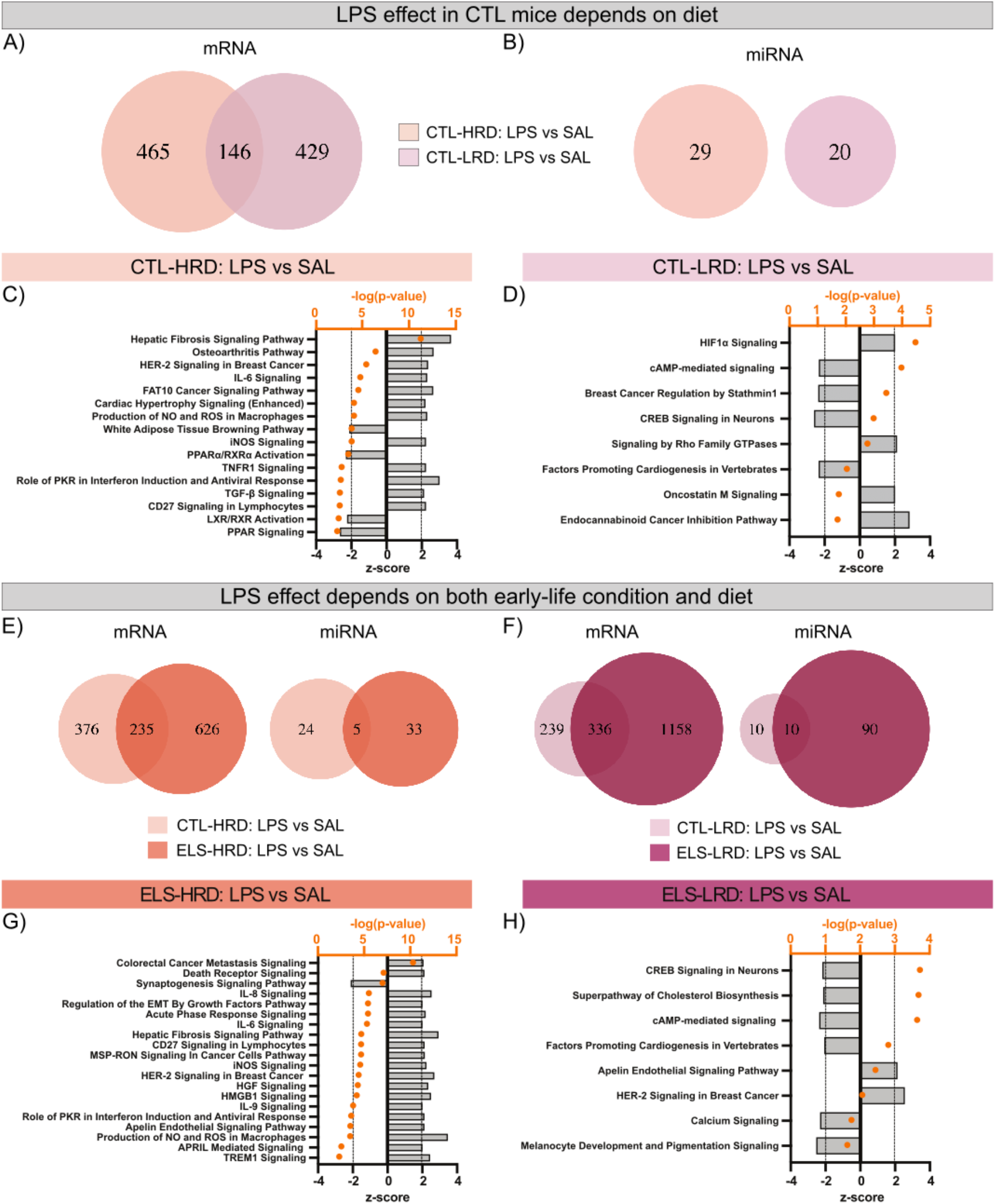
Effects of an acute LPS challenge on hippocampal miRNA and target mRNA expression depend on exposure to ELS and early dietary ω6/ω3 ratio. **AB)** Venn diagrams depicting the diet depending effects in control mice on mRNA **(A)** and miRNA **(B)** expression from the integrated miRNA-mRNA analysis. **CD)** IPA showing the top20 molecular pathways (p-value <0.05 and z-score > |2|) involved in the impact of LPS in control mice fed the HRD **(C)** or the LRD **(D). E)** Venn diagrams depicting the condition dependent effects of LPS on mRNA and miRNA expression in mice fed the HRD. **F)** IPA showing the molecular pathways (p-value <0.05 and z-score > |2|) involved in the impact of LPS in ELS exposed mice fed the HRD. **G)** Venn diagrams depicting the condition dependent effects of LPS on gene and miRNA expression in mice fed the LRD. **H)** IPA showing the molecular pathways (p-value <0.05 and z-score > |2|) involved in the impact of LPS in ELS exposed mice fed the LRD. Abbreviations: CTL: control, ELS: early-life stress, SAL: saline, LPS: lipopolysaccharide, HRD: high ω6/ω3 ratio diet, LRD: low ω6/ω3 ratio diet.

Next, we investigated whether ELS impacts the LPS response and if this depends on the early diet by comparing mRNA and miRNA expression profiles in response to LPS in CTL and ELS exposed mice under both dietary conditions. First, in mice fed the HRD (*CTL-HRD: LPS versus SAL* as compared to *ELS-HRD: LPS versus SAL*), the integrated miRNA-mRNA analysis detected 614 mRNAs regulated by 29 miRNAs (as mentioned also above). For ELS mice fed the HRD, LPS induced 868 DEGs regulated by 38 miRNAs. The majority of mRNAs and miRNAs were unique for either control or ELS exposed mice (**Fig. 3E**). Comparable to the miRNAs in CTL mice, also for ELS mice fed the HRD most miRNAs were upregulated by LPS (95%; e.g. miR-27b-5p, miR-138-2-3p, miR-187-3p, miR-138-5p, miR-181a-5p, miR-146a-5p). IPA on the target genes reveal that both lists include inflammatory/immune-related signaling pathways such as “IL-6 signaling”, “Production of Nitric Oxide and Reactive Oxygen Species in Macrophages”. While there were also condition dependent pathways, such as for CTL mice specifically “PPARα/RXRα activation” and “TGFβ signaling” and for ELS mice inhibition of “synaptogenesis signaling” and “TREM1” signaling (**Fig 3F; supplementary table S6**).

Second, we investigated the impact of ELS on the LPS response in mice fed the LRD (*CTL-LRD: LPS versus SAL* as compared to *ELS-LRD: LPS versus SAL*). As noted above, integration analysis for the LPS response in CTL mice fed the LRD detected 580 mRNAs regulated by 20 miRNAs. For ELS exposed mice a substantially higher number of miRNAs and target mRNAs were detected, 1502 mRNAs regulated by 100 miRNAs Also here, most of the differentially expressed mRNAs and miRNAs were unique for either control or ELS exposed mice (**Fig. 3G**). Similar to CTL mice fed the LRD, also in ELS mice fed the LRD the majority of miRNAs were downregulated by LPS (90%; e.g. miR-30b-5p, miR-30c-1-3p, miR-181d-5p, miR-30e-3p, miR-138-2-3p, miR-99b-5p, miR-99a-5p, miR-149-5p, miR-187-3p, miR-378a-3p, miR-138-5p, miR-221-3p, miR-26b-5p, miR-125a-5p, miR-125b-5p, miR-27b-3p, miR-145a-5p, miR-30b-3p), which was even more pronounced as compared to CTL mice (**Table 1; supplementary table S5**). IPA analysis revealed that in both CTL and ELS mice fed the LRD the pathways “cAMP-mediated signaling” and “CREB signaling in neurons” were inhibited in response to LPS, while the other affected pathways were dependent on condition (CTL/ELS). For CTL mice specifically we detected for example activation of “Signaling by Rho Family GTPases” and “Endocannabinoid Cancer Inhibition Pathway” and for ELS mice specifically inhibition of for example “Superpathway of Cholesterol Biosynthesis” and “Calcium signaling” (**Fig. 3H; supplementary table S6**).

In summary, as expected there is a strong hippocampal gene and miRNA expression response to an acute LPS challenge under all conditions, however both condition and diet lead to unique alterations in miRNAs, target mRNAs and associated molecular pathways. In both CTL and ELS exposed mice fed the HRD most pathways were activated and associated with inflammation, while in mice fed the LRD this was not the case, rather an inhibition was detected in pathways associated with hippocampal plasticity. ELS led to increased numbers of differentially expressed miRNAs and target mRNAs and differential activation and inhibition of the associated pathways.

## 4. Discussion

In this study, first we confirm our previously reported protective effect of the low ω6/ω3 diet from postnatal day (P)2 to P42 against ELS-induced cognitive deficits^20^. Furthermore, by using an integrated analysis of genome wide miRNAs and mRNAs profile, we show that: i) early dietary ω6/ω3 PUFA ratio has long-term effects on hippocampal miRNA and target mRNA expression and determines the specific impact of ELS, leading to entirely different profiles depending on the diet, ii) the ELS induced cognitive impairments in mice fed the high ω6/ω3 diet seem to be mediated by a reduction in hippocampal miRNA expression leading to altered target mRNA expression associated with protein kinase A (PKA) and ephrin signaling, iii) ELS mice fed the low ω6/ω3 diet exhibit altered miRNA and target mRNA expression (as compared to CTL mice fed the same diet) ultimately leading to an activation of pathways associated with hippocampal plasticity and learning and memory (e.g. CREB signaling), providing molecular substrates underlying the beneficial effects of the LRD on ELS induced cognitive impairments and iv) finally we demonstrate that both ELS and early diet determine the LPS-induced alterations in the miRNA and mRNA expression profile and associated molecular pathways. For example, in mice exposed to the high ω6/ω3 diet we detected mostly an activation of pathways associated with inflammatory signaling, while in mice fed low ω6/ω3 diet this was significantly reduced and rather an inhibition was found in pathways associated with hippocampal plasticity, demonstrating both an anti-inflammatory property of the diet as well as its involvement in hippocampal plasticity.

### 4.1 Molecular mechanisms underlying ELS-induced cognitive deficits and those of the beneficial effects of the PUFA diet

We confirm the previously reported ELS induced cognitive impairments in adult mice fed the high ω6/ω3 diet early in life and the rescue by the low ω6/ω3 diet from P2 until P42^20^. Our results are also in line with the evidence supporting that ω3 fatty acids are important for cognitive functioning in rodents^37,38,68,69^, and that specifically adequate ω3 fatty acid availability during early sensitive stages of development is critical for later life cognitive outcome^26,70^. Our data supports the notion that a relatively short and subtle modulation of the ω6/ω3 ratio during early-life can protect against ELS-induced cognitive deficits in adulthood.

#### 4.1.1 Long-term effect of the early dietary PUFAs on hippocampal miRNAs and mRNAs profile

Firstly, we briefly discuss the effects of the diet under basal conditions in control mice. Unsupervised WGCNA identified a diet associated gene co-expression module that was related to protein phosphorylation and nervous system development. Genes with highest centrality in this module were for example Efna3 (or Ephrin A3) and Tumor Necrosis Factor Receptor Superfamily Member 9 (Tnfrsf21 – also known as death receptor 6 (DR6)). Both genes were previously associated with neuronal plasticity processes; i.e. Efna3 with neuronal differentiation and synaptic plasticity (long-term potentiation)^71,72^, and Tnfrsf21 with axonal pruning both in the developing and aging brain^73,74^. Also, our integrated analysis of mRNA and miRNA expression suggests that under basal conditions there are long-term effects of early dietary PUFAs on miRNAs and target mRNAs related to hippocampal plasticity. Our data is in line with other studies investigating ω3 PUFA supplementation and hippocampal function and learning and memory^75,76^. Importantly, our dietary intervention ended months prior to RNA analysis, pointing to long-lasting programming effects of early dietary PUFAs on miRNAs, mRNAs and associated molecular pathways. In particular, our combined miRNA-mRNA analysis revealed that some of the miRNAs altered by the diet, such as miR-381-3p and miR-200c-3p, were previously linked to neuronal plasticity processes^77^ and several target mRNAs were associated “cAMP-response element binding protein (CREB)”. These include Metabotropic Glutamate Receptor 4 (GRM4), key for synaptic transmission and hippocampus-dependent learning^78^, and bone morphogenetic protein 1a (BMPR1A) which plays a role in hippocampal neurogenesis and fear behaviours^79^. Others have shown more direct modulation of cAMP/CREB signaling by dietary PUFAs. For example, ω3 PUFA deprivation in rats reduced CREB activity together with BDNF and MAPK activity^80^ and in primates dietary PUFAs were shown to activate CREB via activation of G-protein-coupled receptor 40 (GPR40)^81^. CREB is a transcription factor transmitting extracellular signals to regulate the expression of many genes, including brain-derived neurotrophic factor (BDNF) via which it can increase cell survival^82^. Additionally, the low ω6/ω3 diet increased several miRNAs previously reported to have anti-inflammatory properties, also in the context of microglial cells, e.g. miR-30b-5p, miR-7a-5p, miR-27b-3p, miR-29c-3p, miR-9-5p, miR-30a-5p^39,83^. This data suggests that early dietary PUFAs can have a long-lasting impact on inflammatory regulation via miRNAs. Affected mRNA expression was additionally associated with “phagosome formation”, thereby possibly affecting phagocytosis. Notably, we have previously reported long lasting effects of the early dietary ω6/ω3 ratio on microglial CD68 expression, a marker for phagocytosis. In line with this, studies have shown the ability of dietary PUFAs, i.e. ω3 PUFAs DHA and EPA and their lipid mediators, to modulate microglial phagocytosis of synaptic elements^84^ and of amyloid-β in the context of Alzheimer’s disease mouse model^85,86^. The current data further supports the notion that dietary ω3 PUFA’s, even when only supplied early in life, can lead to changes in molecular pathways associated with phagocytosis into adulthood.

In summary, these observations demonstrate that the early dietary ω6/ω3 ratio affects miRNA and target mRNAs at basal state, associated with inflammatory processes, neuronal plasticity and cell-cell communication via affecting CREB/cAMP signaling and phagosome formation. The current data is further evidence for long-lasting effects of early dietary PUFAs on later-life molecular pathways thereby possibly affecting neuronal functions.

#### 4.1.2 Impact of ELS on integrated hippocampal miRNA and gene expression profile depends on early dietary PUFAs

ELS leads to cognitive impairments in mice fed a standard diet with a high ω6/ω3 ratio^65^, which makes it key to understand which molecular substrates could underlie these cognitive deficits. The unsupervised WGCNA detected ELS mediated co-expression of genes associated with structural and functional components of the synapse and axon guidance/axonogenesis, supporting the notion that these processes are modulated by ELS and might underlie ELS induced cognitive impairments. Overall we detected an ELS mediated downregulation of miRNAs, such as for example miR-200c-3p, miR-182-5p and miR-183-5p, which were previously associated with neuronal plasticity processes^77,87^. In particular, high levels of miR-183 were reported to support long-term memory formation dependent on protein phosphatase 1^88^, suggesting that the observed ELS induced reduction in miR-183-5p expression in mice fed the HRD might play a role in the cognitive deficits seen in these mice. Pathway analysis on the target mRNAs, including guanine nucleotide binding protein Subunit Alpha 13 (GNA13) and tyrosine-protein phosphatase non-receptor type 13 and 14 (PTNPN13/14), revealed a role for Protein Kinase A (PKA) and Ephrin Receptor signaling in the ELS induced deficits. Generally, protein phosphatases are believed to be memory suppressors while protein kinases rather support memory formation^89–91^. Eph receptors and their Ephrin ligands can guide axons and induce cellular events that underlie changes in synaptic efficacy^92^. Therefore, they play central roles in memory formation and dysregulation of Ephrin signaling has been reported in diseases that include memory impairments such as Alzheimer’s disease and anxiety-related disorders^93^. The above mentioned molecular pathways might contribute to the ELS-induced alterations in neuronal and synaptic plasticity, such as reduced hippocampal volume, reduced adult neurogenesis, altered neuronal excitability and hippocampal dependent memory deficits in adulthood^4,20,94^. Notably, as mentioned above, WGCNA revealed a diet module related to protein phosphorylation and included hub-gene Efna3, indicating that ELS and diet interact on converging molecular pathways thereby influencing hippocampal plasticity.

Indeed, we find that the effects of ELS on hippocampal mRNA and miRNA expression depends on early dietary ω6/ω3 ratio, with generally more ELS regulated miRNAs and mRNA in mice fed the low ω6/ω3 ratio diet. In ELS mice fed the beneficial low ω6/ω3 diet, as compared to CTL mice fed the same diet, the altered miRNA and mRNAs led to increased activity of “CREB signaling in neurons” and other pathways involved in hippocampal plasticity such as “Endocannabinoid Neuronal Synapse Pathway” and “Calcium signaling”. As mentioned earlier, CREB signaling was also affected, but not significantly activated, by the low ω6/ω3 diet in CTL mice. Thus, early dietary ω6/ω3 ratio, while altering several miRNAs and target mRNAs associated with hippocampal plasticity in CTL mice, it has more pronounced effects in mice previously exposed to ELS, possibly underlying the diet mediated rescue of ELS induced effects on measures of hippocampal plasticity and learning and memory^20^. In addition, specifically in ELS mice fed the low ω6/ω3 diet the pathway “IL15 production” was significantly activated. IL15 has been previously proposed to be a “neuroprotective” cerebral cytokine related to neuronal plasticity by modulating GABAergic transmission and reducing neuronal cell death^95^ and was reported to prevent neuropsychiatric-like symptoms in mice, implicating potential therapeutic role for this cytokine^96^. Supporting this pathway might therefore also be one of the ways via which the low ω6/ω3 diet unravels its beneficial effects on hippocampal functions in ELS exposed mice.

In summary, reduced miRNA expression (e.g., miR-183-5p) and altered PKA and Ephrin Receptor signaling might underlie ELS induced alterations in hippocampal plasticity and memory impairments. Importantly, ELS mice fed the protective low ω6/ω3 diet, which no longer exhibit cognitive deficits, exhibited a largely different set of differentially expressed miRNAs and mRNA when compared to their respective controls. In particular they displayed increased activation of pathways associated with hippocampal plasticity and learning and memory. This data provides us with new mechanistic insides on which molecular pathways could underlie the beneficial effects of the low ω6/ω3 diet early in life on hippocampal plasticity and cognition in adulthood.

#### 4.1.3 The response to an inflammatory challenge in adulthood depends on early diet and stress

Both WGCNA and the mRNA-miRNA integrated analysis show a strong impact of an acute LPS challenge on genes related to the inflammatory response, disease states and cellular stress. Next to these expected alterations, differences were present in the response to LPS depending on ELS and diet, with the early dietary ω6/ω3 ratio having the largest impact on miRNAs and their target mRNAs.

Regarding the influence of diet, in mice fed the high ω6/ω3 diet the majority of miRNAs were upregulated by LPS and mRNAs were mostly associated with the inflammatory response and cellular stress. This points to miRNA mediated regulation of genes involved in the inhibition of the inflammatory response which indeed for specific miRNAs has been reported earlier^83^. In mice fed the low ω6/ω3 diet however, we detected an entirely different miRNA and mRNA profile in response to LPS. By far the majority of miRNAs were downregulated by LPS and target mRNAs were mostly associated with hippocampal plasticity such as “cAMP-mediated signaling” and “CREB signaling in neurons”, similar to as the pathways affected by the diet under basal conditions. However, while under basal conditions we detected a low ω6/ω3 diet specific activation of these pathways in particular in mice previously exposed to ELS, in response to LPS hippocampal plasticity pathways were downregulated in both control and ELS exposed mice fed the low ω6/ω3 diet. While no studies have investigated the effect of early dietary PUFAs on genome wide gene expression in response to LPS, there is evidence that LPS induces expression of inflammatory genes in the hippocampus, while it inhibits genes associated with learning and memory^97^. In the current study, this effect was most pronounced in ELS exposed mice fed an early diet with low ω6/ω3 PUFA ratio, supporting the notion that these pathways were activated by the diet under basal conditions.

Next, also ELS impacted the LPS induced alterations in miRNA and target gene expression. Firstly, ELS increased the amount of differentially expressed miRNAs and mRNAs in response to LPS as compared to CTL mice, leading to differential activation of inflammatory pathways. This is in line with previous work showing that stress exposure early in life modulates the adult response to inflammatory challenges^2,13–15,17^. When we look specifically at the LPS inhibited pathways, we detected condition dependent changes that were mostly related to processes associated with hippocampal plasticity, supporting our finding that these pathways were differentially activated between CTL and ELS mice under basal conditions. The current data demonstrates early programming of the hippocampus by ELS leading to differential gene expression response to an inflammatory challenge in adulthood mediated by altered miRNAs. It would be of interest to know whether the miRNA changes are already present early in life and which overlap with plasma miRNA. In fact, we have previously reported on the potential of plasma miRNAs from human adults exposed to ELS as predictive biomarkers for later life mental disorders^48,49^.

In summary, our integrated analysis of miRNAs and their mRNAs shows a strong impact of the early environment (ELS and diet) on the effect of LPS. In mice fed a high ω6/ω3 diet LPS increases miRNA expression and activates pathways associated with inflammation and cellular stress, while in mice fed the low ω6/ω3 diet LPS reduces miRNA expression, reduces activation of inflammatory pathways and specifically inhibits pathways associated with hippocampal plasticity and learning and memory. Moreover, the high ω6/ω3 diet specific activation of inflammatory pathways and low ω6/ω3 diet inhibition of hippocampal plasticity pathways were more pronounced in mice previously exposed to ELS, demonstrating its long-term modulation of miRNA and mRNA expression response to a later-life inflammatory challenge.

Concluding, by using an integrated approach of combining mRNA and miRNA expression data we show that exposure to stress and dietary PUFA’s early in life has long-lasting programming effects on hippocampal miRNA and target mRNA expression and associated molecular pathways, both under basal conditions and in response to an inflammatory challenge in adulthood. miRNAs clearly play a key role in driving gene expression changes, which depend on the early environment. We provide new molecular insights as to how a low ω6/ω3 diet during development, leading to more ω3 PUFA availability, could exert its long-lasting beneficial effects on hippocampal plasticity and learning and memory especially in vulnerable populations exposed to stress and/or inflammation.

## Author contributions

KR and MRA conceptualized this study and performed all mouse-related experimental work. KR and NL analyzed the microarray data with additional help of HJE regarding the WGCNA. KR prepared all figures and tables and wrote the manuscript. AK conceptualized and supervised this study and reviewed the manuscript. All authors contributed to editing the manuscript.

